# Conflicting Bottom-up and Top-down Signals during Misrecognition of Visual Objects

**DOI:** 10.1101/521252

**Authors:** Mohamed Abdelhack, Yukiyasu Kamitani

**Affiliations:** Neuroinformatics Laboratory, Department of Intelligence Science and Technology, Graduate School of Informatics, Kyoto University, Kyoto, Japan; Computational Neuroscience Laboratories, ATR, Kyoto, Japan; Department of Psychiatry, Graduate School of Medicine, Kyoto University, Kyoto, Japan

**Keywords:** top-down, bottom-up, hierarchical, visual processing, neural representation enhancement, deep neural networks, blurred image, misrecognition

## Abstract

Visual recognition involves integrating visual information with other sensory information and prior knowledge. In accord with Bayesian inference under conditions of unreliable visual input, the brain relies on the prior as a source of information to achieve the inference process. This drives a top-down process to improve the neural representation of visual input. However, the extent to which non-stimulus-driven top-down information affects processing in the ventral stream is still unclear. We conducted a perceptual decision-making task using blurred images, while conducting functional magnetic resonance imaging. We then transformed brain activity into deep neural network features to distinguish bottom-up and top-down signals. We found that top-down information unrelated to the stimulus had a minimal effect on lower-level visual processes. The neural representations of degraded stimuli that were misrecognized were still correlated with the correct object category in the lower levels of processing. In contrast, activity in the higher cognitive areas was more strongly correlated with recognition reported by the subjects. The results indicated a discrepancy between the results of processing at the lower and higher levels, indicating the existence of a stimulus-independent top-down signal flowing back down the hierarchy. These findings suggest that integration between bottom-up and top-down information takes the form of competing evidence in higher visual areas between prior-driven top-down and stimulus-driven bottom-up signals. These findings could provide important insight into the different modes of integration of neural signals in the visual cortex that contribute to the visual inference process.

## 1 Introduction

Natural visual recognition relies not only on the processing of visual input, but also on other sensory, contextual, memory, and attentional cues, particularly under visually challenging conditions. Recognition is attained through an integration of bottom-up, recurrent, and top-down pathways that is believed to follow a Bayesian process (Lee and Mumford, 2003; Heeger, 2017; Beck et al., 2008; Aitchison and Lengyel, 2017). Several previous studies have investigated various mechanisms of integration when degraded sensory input is introduced. Top-down signals were reported to cause bias in the early visual and motion sensitive areas as a result of presenting a prior (Kok et al., 2013, 2012). Other studies reported a sharpening effect of top-down signals for improving neural representations due to degraded image viewing (Abdelhack and Kamitani, 2018; Hsieh et al., 2010). Although these studies addressed the question of top-down modulation of neural representations due to visual stimuli, they failed to explain the neural dynamics at play when an incoming stimulus is misrecognized. In the current study, we introduce a novel mode of integration between signals in the visual system where bottom-up and top-down signals supply conflicting information, causing misrecognition when the input stimulus is unreliable.

To investigate the different levels of representation of neural signals and top-down effects in visual and cognitive systems, we use a deep neural network (DNN) representational space. DNN models have been shown to provide a good proxy for understanding these top-down effects in the visual system, because the models typically used for object recognition consist only of a feedforward (bottom-up) processing pathway. Moreover, DNN models have been shown to follow a similar hierarchy to that in the primate visual system (Horikawa and Kamitani, 2017; Cadieu et al., 2014; Khaligh-Razavi and Kriegeskorte, 2014; Yamins et al., 2014; Güçlü and van Gerven, 2015). DNNs have also been utilized to demonstrate top-down sharpening of neural representations of blurred images in the human brain (Abdelhack and Kamitani, 2018).

In the current study, we utilize DNN models as a proxy for neural activation from functional magnetic resonance imaging (fMRI) to identify bottom-up signals as features of the true category of a blurred image, and top-down signals as the features of the category identified by the subject during an image recognition task. Under conditions of misrecognition, the results reveal that the bottom-up and top-down signals are incongruent. We propose the existence of an integration point at which the reported category (during both correct and incorrect recognition) is represented. Thus, the current findings indicate the existence of Bayesian-like neural computation in the visual system. The results shed new light on the interaction between the bottom-up and top-down pathways. These findings could be beneficial in the investigation of circular inference that occurs in psychotic conditions like schizophrenia (Jardri et al., 2017).

## 2 Materials and Methods

The data used in this study were obtained from a previous study in our laboratory (Abdelhack and Kamitani, 2018). Here we provide a brief description of the experimental methods, focusing on differences in data processing and analysis. For a complete account please refer to our previous study (Abdelhack and Kamitani, 2018).

### 2.1 Subjects

We conducted fMRI experiments with five healthy subjects (three males and two females; age range: 22 to 33 years) with normal or corrected-to-normal vision. The study protocol was approved by the Ethics Committee of ATR. All the subjects provided written informed consent before participation in the experiments.

### 2.2 Visual stimuli

Image stimuli were obtained from the ImageNet online database (Jia Deng et al., 2009), and were initially used for training, testing, and validation of the DNN model employed in our study (see below). In this study, we presented both blurred and original images. For blurred image stimuli, three levels of blur were used. A square-shaped averaging filter was applied to each image. The size of the filter (x) relative to the image size (500 pixels) dictates the degradation level. We selected three different blur levels for our protocol (x/500 = 25%, x/500 = 12%, and x/500 = 6%). Larger filters correspond to more blurring of the image.

### 2.3 Experimental design

We conducted two experimental runs, one for decoder training where we presented natural undegraded image stimuli to the subjects, and another test experiment in which we presented blurred image stimuli. In decoder training runs, 1,000 images were shown to the subject, and each image represented one category from the AlexNet classification categories as detailed in our previous study (Abdelhack and Kamitani, 2018). In the test image runs, subjects were shown images belonging to one of five categories (airplane, bird, car, cat, or dog). We presented the stimuli in smaller sequences starting with the most blurred image for one stimulus, and the blur level gradually decreased until reaching the original image (25%, 12%, 6%, and 0% blurring; Figure 1A). We selected this order of presentation to avoid subjects having a memory-prior of the less blurred stimuli when viewing the more blurred images, and to investigate the evolution of correct recognition of the stimulus. We presented 20 mini-sequences (four images each) randomized over two runs (40 stimulus images each). Images were all presented in the center of the screen with a size of 12 × 12° of visual angle using Psychtoolbox (Kleiner et al., 2007) in a flashing sequence for 8 s at 1 Hz (500 ms on-time). For test runs, subjects reported their best guess of the perceived content of the stimulus out of the five categories via voice feedback into an optical microphone placed near the mouth to decrease motion artefacts. Subjects also reported the certainty level of their response by pressing one of two buttons (one for certain and one for uncertain).

**Figure 1.**
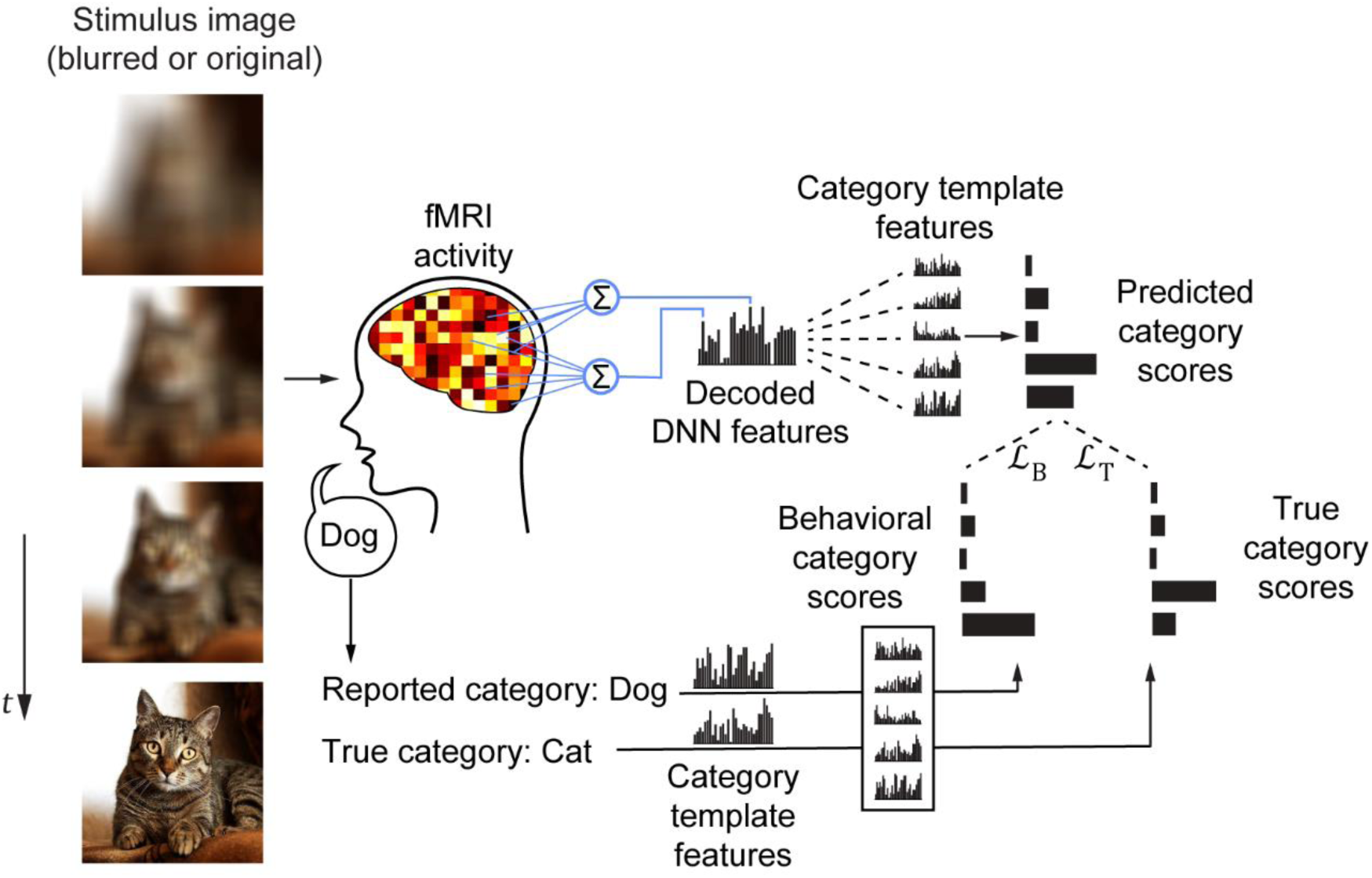
Experimental procedure and main analysis: Each subject viewed blurred images of different blur levels varying from highly blurred to non-blurred. Subjects reported the perceived object category that was used along with the true object category to generate simulated category scores based on behavioral responses and ground-truth. Brain activity recorded while the subject viewed blurred images was used to decode DNN features that were later used to derive the predicted category scores.

### 2.4 MRI data preprocessing

The first 8 s of each acquisition were rejected to avoid scanner instability effects. We then performed preprocessing using SPM8^1^(RRID: SCR_007037) that comprised 3D motion correction, slice-timing correction, and co-registration to the appropriate high-resolution anatomical images and to the T1 anatomical image. We then interpolated the echo planar imaging (EPI) data to 2 × 2 × 2 mm voxels. We further preprocessed the data to prepare it for the decoding algorithm using Brain Decoder Toolbox 2^2^(RRID: SCR_013150). We shifted volumes by 4 s (two volumes) to compensate for hemodynamic delays, then removed the linear trend from each run and normalized the data by subtracting the mean of each voxel time sequence for each run. We then averaged the volumes for each stimulus image (four volumes) to increase the signal-to-noise ratio. These averaged voxel values for each stimulus block were used to construct the input feature vector for the DNN feature decoding analysis.

### 2.5 Region of interest construction

We constructed regions of interest (ROIs) for several regions along the ventral stream, including the lower visual areas V1, V2, and V3, the intermediate area V4, higher visual areas including the lateral occipital complex (LOC), parahippocampal place area (PPA), and fusiform face area (FFA), and a higher cognitive area, the middle temporal gyrus (MTG).

We specified the areas of V1–4 via a retinotopy experiment following a standard protocol (Engel et al., 1994; Sereno et al., 1995). We then analyzed the brain activity and visualized the results on an inflated brain map using FreeSurfer^3^(RRID: SCR_001847). We manually cut the ROIs for each subject and transformed the resulting ROIs to the preprocessed EPI data space.

To delineate the higher visual areas, we conducted a functional localizer experiment following a standard protocol (Kanwisher et al., 1997; Kourtzi and Kanwisher, 2000; Epstein and Kanwisher, 1998) including presentation of intact and scrambled images of objects, faces, and scenes. We analyzed the resulting brain activity and viewed the results on an inflated brain map using FreeSurfer. Regions showing significant activation for object images compared with scrambled object images located in the lateral occipital cortex and fusiform gyrus were used to delineate the LOC. In addition, regions showing significant activation for face images compared with scrambled face images located in the fusiform gyrus and lateral occipital cortex were used to delineate the FFA. Finally, regions showing significant activation for scene images compared with scrambled scene images located in the parahippocampal area and fusiform gyrus were used to delineate the PPA. By aggregating the ROIs of V1–4, LOC, FFA, and PPA, we constructed the visual cortex (VC) ROI.

To delineate the higher cognitive area, middle temporal gyrus (MTG), we utilized the principal gradient maps from (Margulies et al., 2016). We first registered the maps from FreeSurfer’s fsaverage space to each subject’s space, then masked out all brain areas from the gradient except for the occipital and temporal cortices. Using the masked gradient, we then divided 10 ROIs based on their principal gradient percentile. We then selected the ROI that represented the deepest 10% along the gradient. These ROIs were typically situated in the middle of the temporal cortex along the MTG.

### 2.6 Deep neural network model

Similar to our previous study (Abdelhack and Kamitani, 2018), we used the AlexNet model (Krizhevsky et al., 2017) as a model for hierarchical image presentation. We utilized the Caffe implementation of the network packaged for the MatConvNet tool in MATLAB (Vedaldi and Lenc, 2015). AlexNet comprises eight layers, the first five of which are convolutional while the last three are fully-connected. Early layers process local and low level features like edges and contrast, while deeper layers process more categorical aspects. AlexNet has been shown previously to exhibit similar hierarchical processing to that of the visual cortex (Horikawa and Kamitani, 2017). We extracted 1,000 features from each layer of AlexNet based on their decodability from brain data (Horikawa et al., 2018). We labelled the features from each layer as DNN1–DNN8.

### 2.7 Behavioral data collection

The subjects reported their best guess of the perceived stimulus via an optical microphone. The verbal response was then recorded, the audio was transcribed, and the results were revised with each subject to guarantee their accuracy. In the few cases where the subject failed to provide a verbal response, the stimulus block was labelled incorrect and category scores were equally divided between the five categories (0.2 each). Subjects also reported their certainty level by pressing one of two buttons to indicate whether they were certain or uncertain about their verbal response, but these data were not used in this study.

### 2.8 DNN feature decoding

We trained a decoder to predict each of the extracted DNN features from the voxels in each ROI. We used a sparse linear regression (SLR) algorithm (Tipping, 2001; Bishop, 2006) to create linear models. For each DNN feature, the prediction (decoded feature) from a given period of brain activity is expressed by

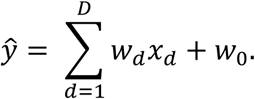

where *x*_*d*_ and *w*_*d*_ are the fMRI signal amplitude and weight of the *d*^th^ voxel in the ROI, *w*_0_ is the bias, and D is the number of the voxels in the ROI. *w*_*d*_ was defined using a training dataset by applying SLR after a voxel selection and normalization procedure (see below).

For each decoder, we conducted voxel selection prior to training by selecting the most strongly correlated 500 voxels with the DNN feature. We then normalized the preprocessed fMRI data for training by scaling them to a mean of zero and a standard deviation of one. The same normalization procedure was also conducted on the target DNN features training dataset. SLR was applied to obtain the weights by iteratively maximizing an objective function (2,000 iterations). SLR involves selecting a small number of voxels as important by estimating the corresponding weights to be nonzero, and ignoring the other voxels by estimating their weights to be zero (Tipping, 2001; Bishop, 2006). For more details on the procedure, refer to Horikawa and Kamitani (2017). When applying the models to test data, fMRI data were normalized using the mean and standard deviation values from the training dataset. The resulting features were extracted as is, and were not denormalized, to avoid any baseline correlations in the subsequent analyses. When features extracted from the DNN were used in the subsequent analyses they were also normalized by subtracting the mean and standard deviation extracted from the training dataset.

For each layer of the DNN, we calculated the decoded feature for each of the DNN units in the layer, and we collectively denoted the decoded features for all units in the layer by a single column vector 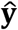 (decoded feature vector).

### 2.9 Category score calculation

To compute decoded and simulated category scores, we first generated template vectors for each category. Similarly to Horikawa and Kamitani (2017), we first used a tool^4^for optimizing an input image to maximize a target category classification score in AlexNet’s last layer, along with a deep generator network that introduced an image prior leading to more natural-appearing images (Nguyen et al., 2016). The deep generator network restricts the feature value space for the lower DNN layers to force them to be more similar to natural images. This assists in making the resultant features relevant in shallower layers as well as deeper layers. AlexNet has 1,000 classification categories as opposed to five in our settings, and our categories cover broader concepts than those of AlexNet. Thus, each of our five categories are covered by several of the classification categories in AlexNet (airplane: two sub-categories, bird: 59 sub-categories, car: 18 sub-categories, cat: five sub-categories, dog: 117 sub-categories). Thus, we generated preferred input images for each of the sub-categories in AlexNet. From those preferred input images, we were able to extract our 1,000 DNN features from each layer and normalize them similar to the decoded features. Then, for each of our five categories, we averaged the features of all corresponding sub-categories to generate one category template feature vector (**z**_c_).

We then calculated decoded category scores for each sample by computing Pearson’s correlation coefficients between this sample’s decoded feature vector 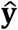 and each category template vector **z**_c_, then normalizing correlation coefficients over all categories using the softmax operation. To compute the simulated category scores for the same sample, we applied the previous operation but in the place of the sample’s decoded feature vector 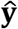, we used one category template feature vector **z**_c_ based on the behaviorally chosen category (behavioral category scores) and the ground-truth category (true category scores).

### 2.10 Analyzing the similarity between decoded and simulated category scores

To analyze the extent to which neural signals in one ROI mimic each of the behavioral response outcomes and ground-truth data, we computed the similarity between the decoded category scores and behavioral category scores (𝓛_B_) and the similarity between decoded category scores and true category scores (𝓛_T_). We first concatenated category scores (both decoded and simulated) for different samples to form vectors, then computed Pearson’s correlation coefficients to represent similarity values (𝓛_B_ and 𝓛_T_). We concatenated all samples (Figure 2) leading to many similarities between 𝓛_B_ and 𝓛_T_ due to the prevalence of correctly recognized samples. We then only concatenated misrecognized samples pooling different blur levels (Figure 3) to isolate samples in which 𝓛_B_ and 𝓛_T_ are incongruent. We finally concatenated the misrecognized samples from the 25% blur levels (Figure 4), as described in section 2.11.

**Figure 2.**
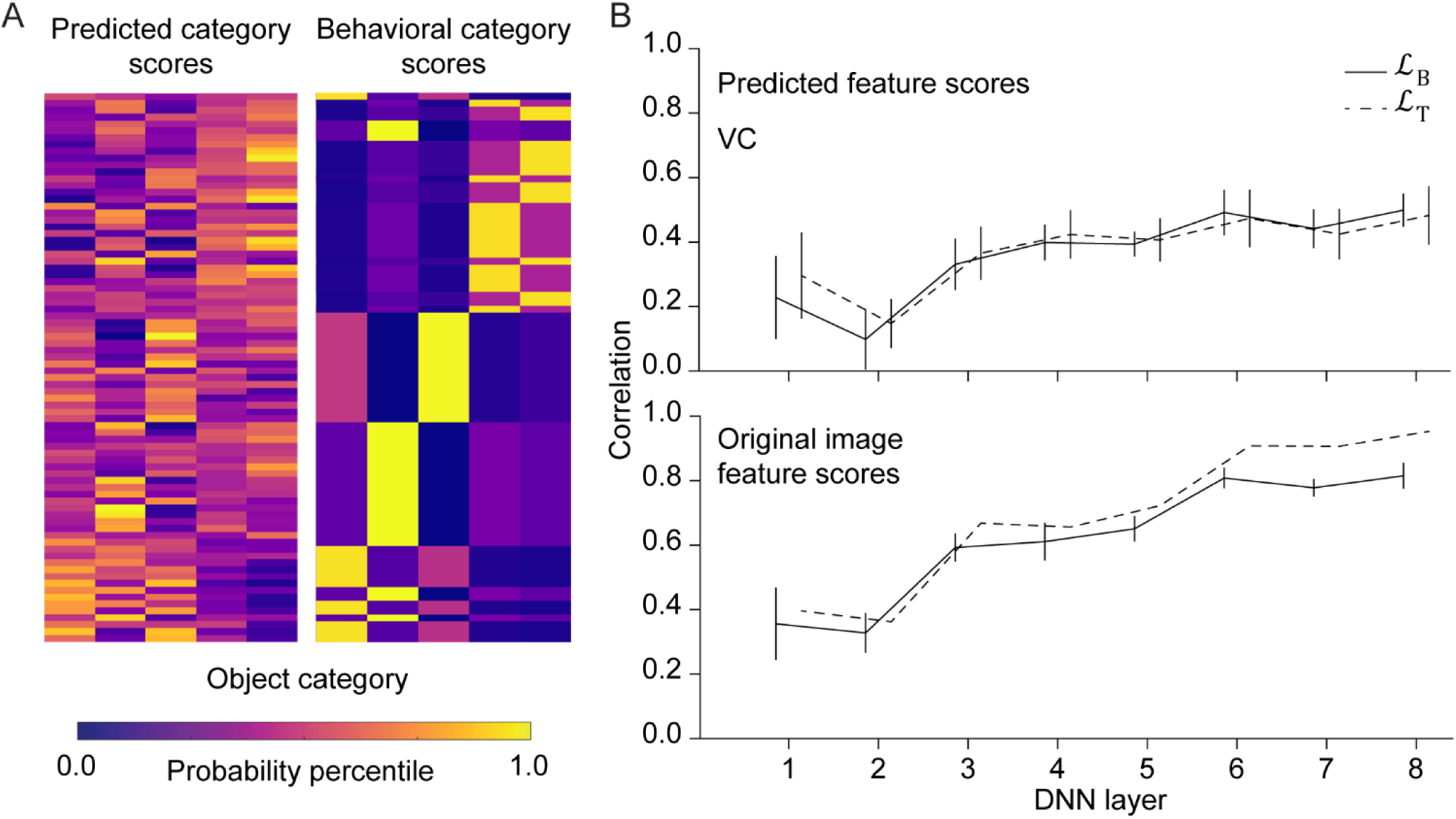
Category scores similarity computation: **A.** Sample of the category scores decoded from brain activity from VC and DNN layer 8 and the corresponding simulated scores based on the behavioral response from subject 1 arranged in a sample by category array. Data are represented as the values of percentiles for each matrix to compensate for any differences in the ranges of values. **B.** Upper: Similarity computation between the predicted category scores from VC and corresponding simulated category scores derived from behavioral response (𝓛_B_) and ground-truth (𝓛_T_). Lower: The same similarity computation using original non-blurred image true features extracted directly from the DNN to compute category probability instead of decoded features. Line plots represent the mean value across five subjects and error bars represent 95% confidence intervals.

**Figure 3.**
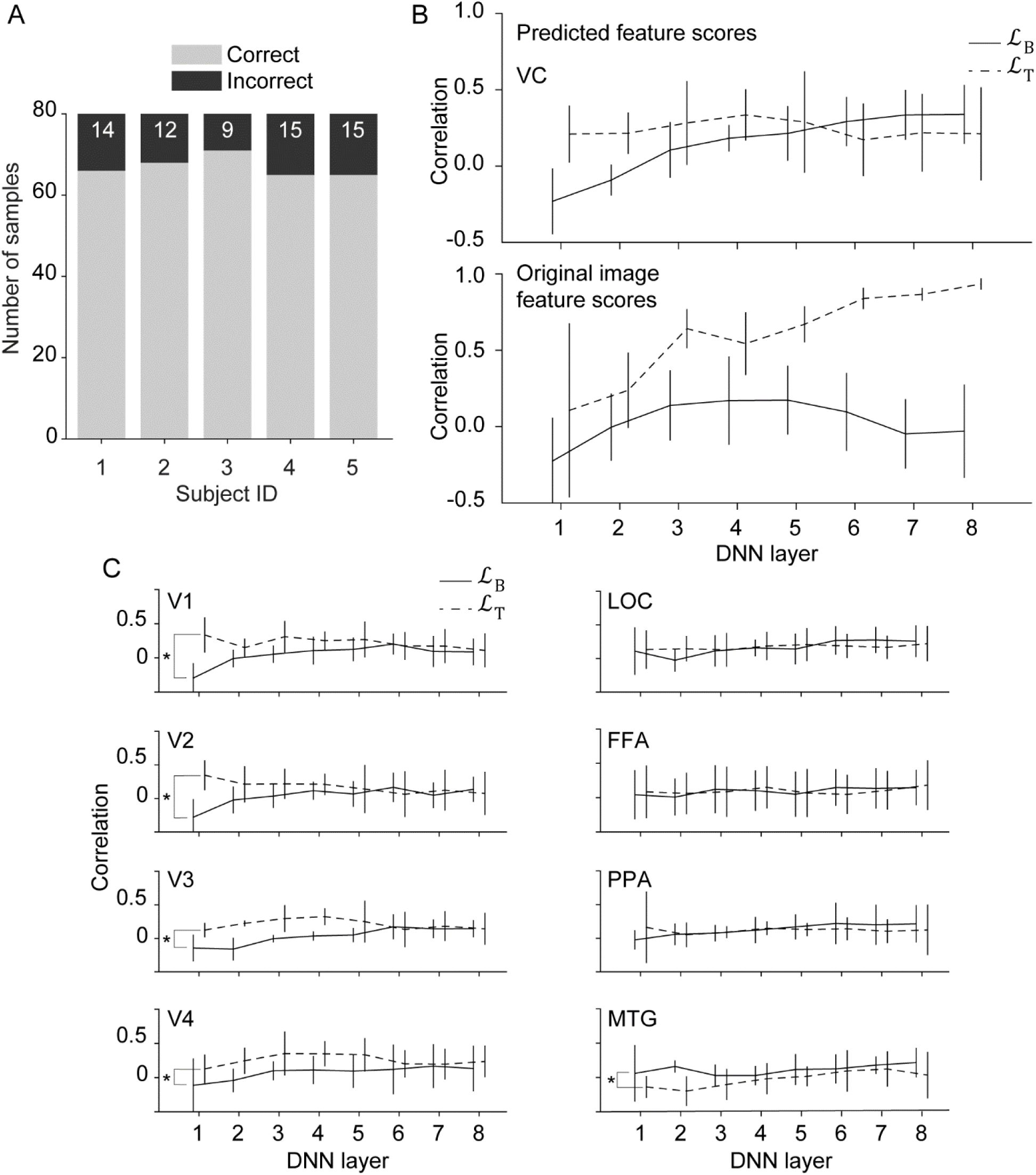
Similarity computation for misrecognized images: (A) A summary of the behavioral response showing the portion of incorrect trials compared with the total number of stimuli for each subject. The number on the incorrect bars indicates the number of incorrect trials. (B) Results of 𝓛_B_ and 𝓛_T_ computation for features predicted from VC for misrecognized trials only (upper) and the same results computed for the original unblurred image features (lower). (C) Results of 𝓛_B_ and 𝓛_T_ computation for features predicted from each ROI for incorrect trials only. Asterisks denote ROIs where 𝓛_B_ and 𝓛_T_ were shown to be significantly different using ANOVA (*p* < 0.005; Bonferroni correction factor = 9). Line plots represent the mean value across five subjects and error bars represent 95% confidence intervals.

**Figure 4.**
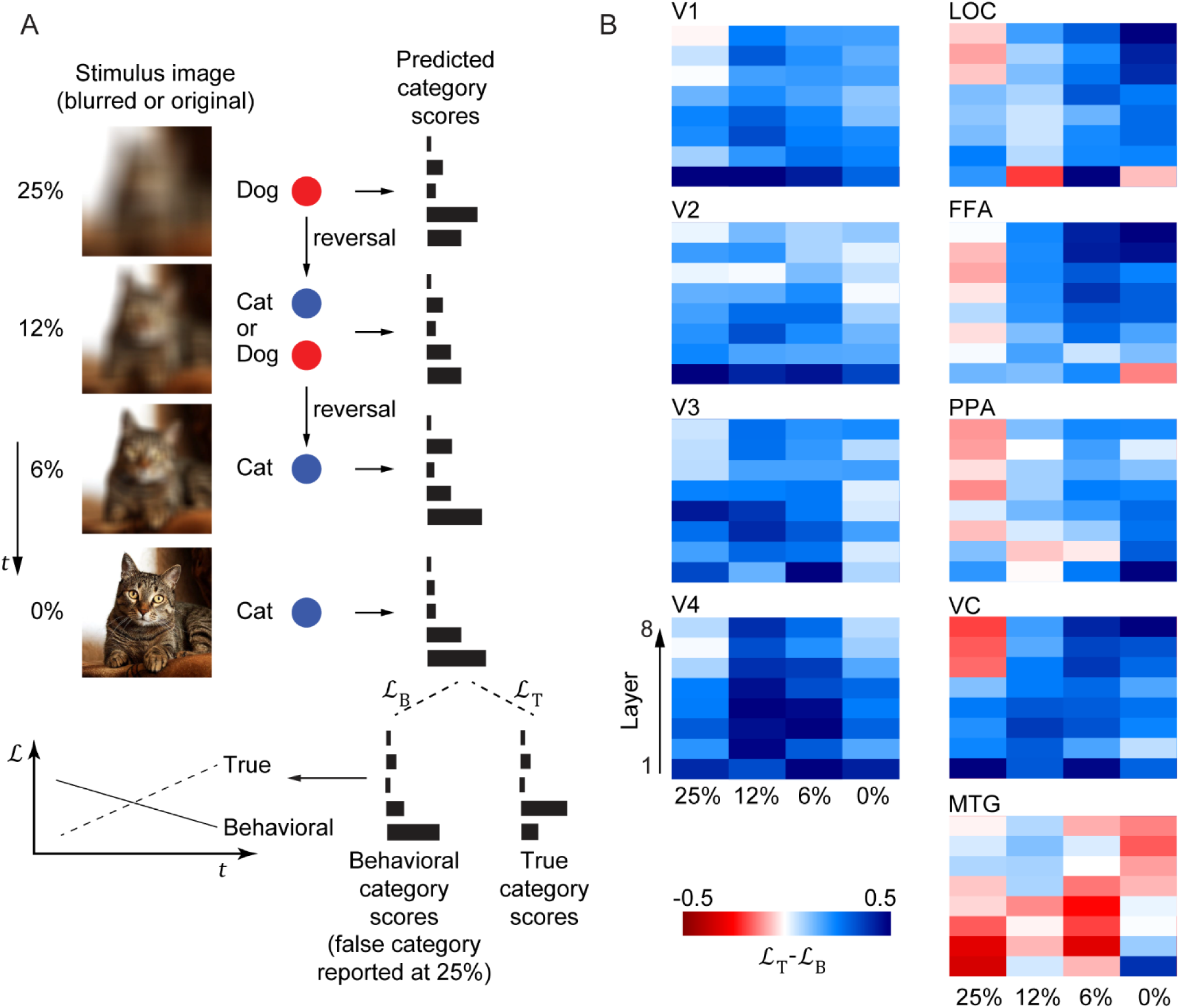
Evolution of correct recognition: (A) Description of the procedure of selecting trials that began with misrecognition followed by an evolution of the response to the correct one as the blur level decreased, showing possible points of recognition reversal. (B) Results of mean 𝓛_T_ − 𝓛_B_ for features predicted from each ROI for different DNN layers as it progresses along the time course of one mini-sequence of incorrect-to-correct recognition.

The similarity values were calculated for each DNN layer features for all five subjects and nine ROIs previously defined. We then calculated the mean similarity across all subjects for each DNN layer and ROI along with the confidence intervals. We also tested the null hypothesis that 𝓛_B_ and 𝓛_T_ originate from the same distribution using analysis of variance (ANOVA) on each ROI, using the DNN layer as an independent variable.

### 2.11 Evolution of correct recognition analysis

To investigate how the correct recognition evolved as the blur level decreased with accumulating evidence, we separated the misrecognized 25% blur level trials and concatenated decoded category scores from those samples. We then obtained decoded category scores from the following trials from each sample’s same mini-sequence until the 0% blur trials irrespective of the behavioral response, and concatenated them in vectors for each blur level. Thus, we obtained decoded category score vectors that represented a temporal evolution from misrecognition (25% level) to correct recognition (0% level). The other two levels comprised a mix of correct and incorrect trials, although predominantly correct trials existed in the two conditions, particularly the less blurred condition (6%). We then concatenated simulated category scores for the 25% blur condition only, to obtain behavioral category scores that were representative of the misrecognition condition and true category scores that represented the correct inference condition (true category scores were actually the same for each blur condition, and did not change across one mini-sequence). We then calculated 𝓛_B_ and 𝓛_T_ for each of the four decoded category score vectors selected earlier with the simulated vectors from the 25% level to visualize how the incorrect category reported at the 25% blur level gets suppressed in favor of the correct category as the blur level decreases (Figure 4A). We then subtracted the similarity values (𝓛_T_ − 𝓛_B_) to obtain a measure of the correct vs. incorrect similarity.

## 3 Results

### 3.1 Calculation of category scores

To compute category scores, we first decoded DNN features from fMRI data of blurred images, similar to the process described in Abdelhack and Kamitani (2018). We then calculated scores from each category for the decoded features using a template matching algorithm (Figure 1). We calculated Pearson’s correlation coefficients between decoded features of each sample and template features of each category (Figure 1). We then normalized the correlation coefficients across all categories using the softmax function to obtain category scores. Moreover, to generate a simulated category probability from the behavioral response and ground-truth data, we applied the same correlation-normalization process, but used the category template feature vectors as input instead of decoded feature vectors. For each sample, the category template vector was selected based on the reported category (behavioral category scores) or the true category (true category scores, Figure 1).

### 3.2 Similarity between decoded and simulated category scores

Using the previous algorithms, we generated category scores for each sample. Figure 2A shows DNN layer 8 decoded category scores from the whole of the visual cortex ROI (VC) of subject 1 arranged in a sample by category array (left panel) and behavioral category scores from the same subject’s behavioral response results (right panel). The two heat maps show similarities in the category score patterns. We then attempted to identify whether the decoded neural behavior corresponded to the behavioral action (top-down) or the ground truth (bottom-up). We concatenated category scores from all samples to produce category score vectors. We estimated how similar the decoded and simulated category scores were by calculating Pearson’s correlation coefficients for each category score vector pair (decoded/behavioral 𝓛_B_ and decoded/ground-truth 𝓛_T_; Figure 1). The resulting similarity signifies which of the bottom-up and top-down signals the decoded category representation mimics. Using DNN features decoded from the VC, we calculated 𝓛_B_ and 𝓛_T_ for all samples (Figure 2B, upper). We observed that the deeper layers showed greater similarity with both the behavioral data and ground truth for both true and predicted features. This is an expected pattern, since earlier layers of the DNN possess topological information that could substantially vary between the template and the stimulus decoded features compared with deeper and more categorical layer features. This was further confirmed when we computed 𝓛_B_ and 𝓛_T_ using category probabilities computed from original non-blurred image true features that were extracted directly from the DNN instead of decoded features.

### 3.3 Similarity patterns in misrecognized trials

One reason for the high correspondence in both conditions is that the high performance of all subjects (Figure 3A) led to many correct trials in which ground-truth and behavioral category scores were exactly the same. To investigate the condition involving discrepancy between bottom-up and top-down signals, we filtered out correctly recognized trials to retain only misrecognized trials and measured the resulting similarity. The results from the VC exhibited an interesting pattern (Figure 3B, upper) in which the earlier DNN layers appeared to show greater similarity with ground-truth category scores (higher 𝓛_T_) as opposed to the deeper layers that showed greater similarity with behavioral category scores (higher 𝓛_B_). These results indicate that while the earlier neural representations point towards the actual image features, further processing leads the brain to misidentify the image as another category. This is in contrast to the original image true features, in which 𝓛_T_ is dominating at all DNN layers, particularly the deeper layers (Figure 3B, lower). These results also indicate that even when the image was misrecognized, the lower-level representations specifying the correct category were still conserved.

When we investigated different ROIs in the visual cortex and higher cognitive areas (Figure 3C), we found that lower and intermediate visual areas exhibited significantly greater similarity with ground-truth category scores (𝓛_T_ > 𝓛_B_; V1: *p* = 4.31 × 10^−6^, V2: *p* = 0.0017, V3: *p* = 1.78 × 10^−7^, V4: *p* = 1.45 × 10^−4^). However, the higher visual areas did not show any significant difference in similarities, although tendencies of 𝓛_B_ > 𝓛_T_ were exhibited in deeper layers. We found that higher cognitive areas located in the MTG exhibited significantly greater similarity with the behavioral category scores (𝓛_B_ > 𝓛_T_; *p* = 0.0011). This ROI was defined from principal gradient analysis (Margulies et al., 2016) as the 90^th^ percentile along the gradient restricting it only to the occipital and temporal lobes of the brain, to capture the ventral stream processing pathway. This area is part of the default mode network, which has traditionally been thought to be active during task negative conditions (Anticevic et al., 2012; Fox et al., 2005) but has more recently been shown to be active during memory retrieval tasks (Sormaz et al., 2018; Konishi et al., 2015; Zhang et al., 2018; Murphy et al., 2018). These results demonstrate that, even in the case of misrecognition, neural representations of the original category are nonetheless conserved in the lower visual areas. These findings also suggest that misrecognition is caused by top-down signals originating from amodal semantic centers.

### 3.4 Evolution of correct recognition

In the previous analysis, we found two opposing neural signals: a stimulus-driven bottom-up signal, and a prior-driven top-down signal. These two pathways integrate and sometimes contradict each other, leading to competing sources of evidence. We investigated how these two signals integrate as the input stimulus evolves through time from unreliable to reliable and recognition evolves from incorrect to correct. We selected only the misrecognized trials from the highest blur level (25%) to create vector decoded category scores. We then created three more predicted category score vectors for the three trials following each of those misrecognized 25% blur, representing the other blur levels (12%, 6%, and 0%) irrespective of the recognition result correctness. We then calculated 𝓛_B_ and 𝓛_T_ using the corresponding simulated category scores solely from the 25% blur level. On the one hand, the ground-truth category score vector did not change across blur levels. On the other hand, the behavioral category score vector varied as recognition changed from incorrect to correct. As we sought to investigate the suppression of the misrecognition, we fixed the behavioral category score vector used in the calculation of 𝓛_B_ over each blur level to the vector of the misrecognized 25% blur level. We then subtracted similarity values (𝓛_T_ − 𝓛_B_) to investigate how the behavioral and ground-truth matrix similarities evolved as the blur level decreased all the way to 0% (Figure 4A). We found that the lower and intermediate visual areas (V1–4) exhibited consistent similarity towards the ground-truth category (Figure 4B). The higher visual areas appeared to exhibit consistent evolution in deeper layers from similarity to the wrong category at 25% and a gradual shift towards the correct category and further increases in similarity as evidence grew stronger. However, the MTG appeared to only exhibit meaningful signals during misrecognition. This behavior may indicate that this region still sends memory-driven signals that get ignored further down the hierarchy due to the relative strength of the stimulus likelihood. Alternatively, this area may cease to function when the bottom-up pathway is more active. These results indicate that the higher visual areas are a likely location for the integration of the bottom-up and top-down signals. These areas may conduct Bayesian estimation using the bottom-up signal as the likelihood signal and the top-down signal as the prior.

## 4 Discussion

In the current study, we investigated the integration of bottom-up and top-down pathways using a blurred image recognition task. Image blurring creates challenging conditions for visual recognition, requiring other sources of information to flow back down the processing hierarchy to reach a conclusion. Integration of the bottom-up and top-down pathways is thought to be conducted in accord with Bayesian inference. We sought to investigate this issue by locating the prior and likelihood signals and the location at which they integrate. We also sought to investigate the extent to which the top-down signals propagate to affect neural representations. We devised an algorithm to calculate semantic category scores represented by brain activity using the DNN as a proxy for neural representations and preferred input images by the DNN units to represent template features of categories presented in the experimental procedure. We then used these category scores to investigate whether different visual and cognitive areas represented the prior or the likelihood signals. To make this distinction, we tested stimuli that exhibited a discrepancy between these two signals. Trials in which subjects failed to correctly identify the category presented in the image were regarded as ones where the likelihood signal reliability is low leading the subject to rely on a contradictory prior signal. Therefore, we examined these misrecognized trials, representing the correct category score as a measure of bottom-up (likelihood) signals, and the incorrectly reported category score as a measure of top-down (prior) signals.

We found that VC ROI activity mimicked the bottom-up signal on the lower processing scales (i.e., the shallower DNN layers) while mimicking the top-down signal on the higher processing levels (Figure 3B). This was further supported by the results in smaller ROIs in the lower and intermediate visual areas (V1–4) that showed a significant preference for the representation of the bottom-up signal (Figure 3C). Higher visual areas exhibited only a tendency for top-down signals, particularly in deeper DNN layers. However, the higher cognitive center in the temporal cortex (the MTG) exhibited a significant preference for the top-down representation, suggesting a role for this region in supplying the prior signals. The MTG is part of the anterior temporal lobe (ATL), that has been suggested as a possible source of top-down signals that influence object recognition (Chiou and Lambon Ralph, 2016). The ATL has also been reported by several studies to function as an amodal semantic hub for sensory signals incoming from different modalities (Chiou and Lambon Ralph, 2018; Levy et al., 2004; Lambon Ralph, 2014; Lambon Ralph et al., 2017; Patterson et al., 2007; Bonner and Price, 2013). In addition, we also constructed the MTG in our study based on its location in the principal gradient (Margulies et al., 2016), meaning that this region is a location at which signals from different modalities converge, and is not modality-bound itself. In addition, the MTG is part of the default mode network that is reported to exhibit activation during memory retrieval tasks (Sormaz et al., 2018; Konishi et al., 2015; Murphy et al., 2018), suggesting that incorrect recognition may emanate from the use of memory as a prior signal when a visual signal is unreliable (Figure 3D). Moreover, although this area is considered semantic (Lambon Ralph et al., 2017), the current results reveal that a significant portion of the lower level information of the prior was represented there (Figure 3C). This notion is in accord with previous reports that show default mode network regions activated during memory retrieval of simple shapes (Murphy et al., 2018; Konishi et al., 2015; Sormaz et al., 2018).

Results from lower visual areas showed also that a top-down prior was unable to propagate all the way to the lower visual areas. Although those areas have been reported to exhibit sharpening behavior (Kok et al., 2012; Abdelhack and Kamitani, 2018; Teufel et al., 2018), the current results indicate that this may not be sufficient to change the original category identity. One possibility is that the subject may find a blurred image too difficult to recognize, leading the brain to give up on sharpening the neural signals and relying on the prior, directly leading to the conservation of representations in the lower visual areas. This may explain the low level of sharpening observed in lower processing levels in highly blurred images (Abdelhack and Kamitani, 2018). Although the higher visual areas did not show significant favorability for any of the bottom-up or top-down signals in misrecognized trials, they showed a slight tendency towards the top-down signals representing the behavioral response, particularly in higher levels of processing (DNN6–8; Figure 3C). Furthermore, when we investigated how recognition evolves from incorrect to correct, activity in the deep layers appeared to follow the behavioral response relatively closely (Figure 4B). This finding may imply that these areas are a point of integration of the prior and likelihood signals to compute the posterior probability, but only semantically, and discarding topological information. The MTG, however, appeared to only exhibit a behaviorally meaningful and relevant prior during incorrect recognition trials. This may indicate that this area was still be providing top-down signals that were ignored at the integration point due to the increasing reliability of the input stimulus. Alternatively, the results could also be explained by the switching observer model (Laquitaine and Gardner, 2018), in which the prior ceases to be provided when the stimulus can be reliably processed. This notion is in accord with the traditional view of the default mode network as a network of task-negative regions (Anticevic et al., 2012; Fox et al., 2005), so, as the input signals become more reliable, the visual cortex engages in the task of visual processing and the default mode network switches away from the task of providing prior signals. The MTG was previously shown to primarily contribute in situations in which input requires supplementation (Chiou and Lambon Ralph, 2016).

We propose a model of visual inference under challenging conditions in which prior signals propagate down from semantic cognition areas while neural responses to visual signals propagate up the visual hierarchy to compete for dominance in the decision making process via Bayesian integration in the higher visual areas in the inferior temporal cortex. In the current study, the prior signal was not explicitly defined and its computation mechanism could vary depending on each subject. It would, however, be more tenable to provide the subjects with explicit prior and investigate cases of conforming and conflicting priors and likelihoods, similar to previous behavioral studies (Jardri et al., 2017). Another limitation of the current study design is that, due to the reliance on fMRI signals, it was difficult to resolve the finer temporal scales of the Bayesian process. Further studies utilizing tools with higher temporal resolution, such as magnetoencephalography, could provide deeper insights into the time progression of each of the prior and likelihood signals along the processing gradient. Nevertheless, the current findings provide insight into the processing of visual and cognitive information in tandem and how different visual processing pathways balance and interact with cognitive pathways to integrate their information. This understanding could provide us with a better understanding of the vision process and with tools to compensate for disruptions in the balance of these pathways that occurs in psychotic conditions, such as autism (Lai et al., 2014) and schizophrenia (Jardri and Denève, 2013; Friston et al., 2016). The current study could also contribute to extending the understanding of conditions that affect cognitive centers, such as semantic dementia, (Patterson et al., 2007) on the visual processing pathways.

## 6 Author Contributions

MA and YK designed the study. MA performed experiments and analyses. MA and YK wrote the paper.

## 8 Acknowledgments

The authors thank Tomoyasu Horikawa, Kei Majima, Mitsuaki Tsukamoto, and Shuntaro Aoki for help with data collection and analysis. We also thank the members of Kamitani Lab for their valuable comments on this manuscript. This research was supported by grants from JSPS KAKENHI Grant numbers JP15H05920, JP15H05710, and ImPACT Program of Council for Science, Technology and Innovation (Cabinet Office, Government of Japan), the Strategic International Cooperative Program (JST/AMED), and the New Energy and Industrial Technology Development Organization (NEDO), and Japanese Government Scholarship (MEXT).

We thank Benjamin Knight, MSc., from Edanz Group (www.edanzediting.com/ac) for editing a draft of this manuscript.

This study was conducted using the MRI scanner and related facilities of Kokoro Research Center, Kyoto University

## 9 Data Availability Statement

FMRI preprocessed data used in this study is available for download from http://www.doi.org/10.6084/m9.figshare.7562516. Codes for analysis are also deposited in the following repository https://github.com/KamitaniLab/BlurMisrecognition.

## Footnotes

1 http://www.fil.ion.ucl.ac.uk/spm,

2 https://github.com/KamitaniLab/BrainDecoderToolbox2,

3 https://surfer.nmr.mgh.harvard.edu/,

4 https://github.com/KamitaniLab/cnnpref,

